# Treatment of advanced atherosclerotic mice with the senolytic agent ABT-263 is associated with reduced indices of plaque stability and increased mortality

**DOI:** 10.1101/2023.07.12.548696

**Authors:** Santosh Karnewar, Vaishnavi Karnewar, Laura S. Shankman, Gary K. Owens

## Abstract

The use of senolytic agents to remove senescent cells from atherosclerotic lesions is controversial. A common limitation of previous studies is the failure to rigorously define the effects of senolytic agent ABT-263 (Navitoclax) on smooth muscle cells (SMC) despite studies claiming that they are the major source of senescent cells. Moreover, there are no studies of the effect of ABT-263 on endothelial cells (EC), which along with SMC comprise 90% of α-SMA^+^ myofibroblast-like cells in the protective fibrous cap. Here we tested the hypothesis that treatment of advanced atherosclerotic mice with the ABT-263 will reduce lesion size and increase plaque stability. SMC (Myh11-CreER^T2^-eYFP) and EC (Cdh5-CreER^T2^-eYFP) lineage tracing *Apoe*^-/-^ mice were fed a WD for 18 weeks, followed by ABT-263 100mg/kg/bw for six weeks or 50mg/kg/bw for nine weeks. ABT-263 treatment did not change lesion size or lumen area of the brachiocephalic artery (BCA). However, ABT-263 treatment reduced SMC by 90% and increased EC-contributions to lesions via EC-to-mesenchymal transition (EndoMT) by 60%. ABT-263 treatment also reduced α-SMA^+^ fibrous cap thickness by 60% and increased mortality by >50%. Contrary to expectations, treatment of WD-fed *Apoe*^-/-^ mice with the senolytic agent ABT-263 resulted in multiple detrimental changes including reduced indices of stability, and increased mortality.

**Graphical abstract:** Uploaded separately.

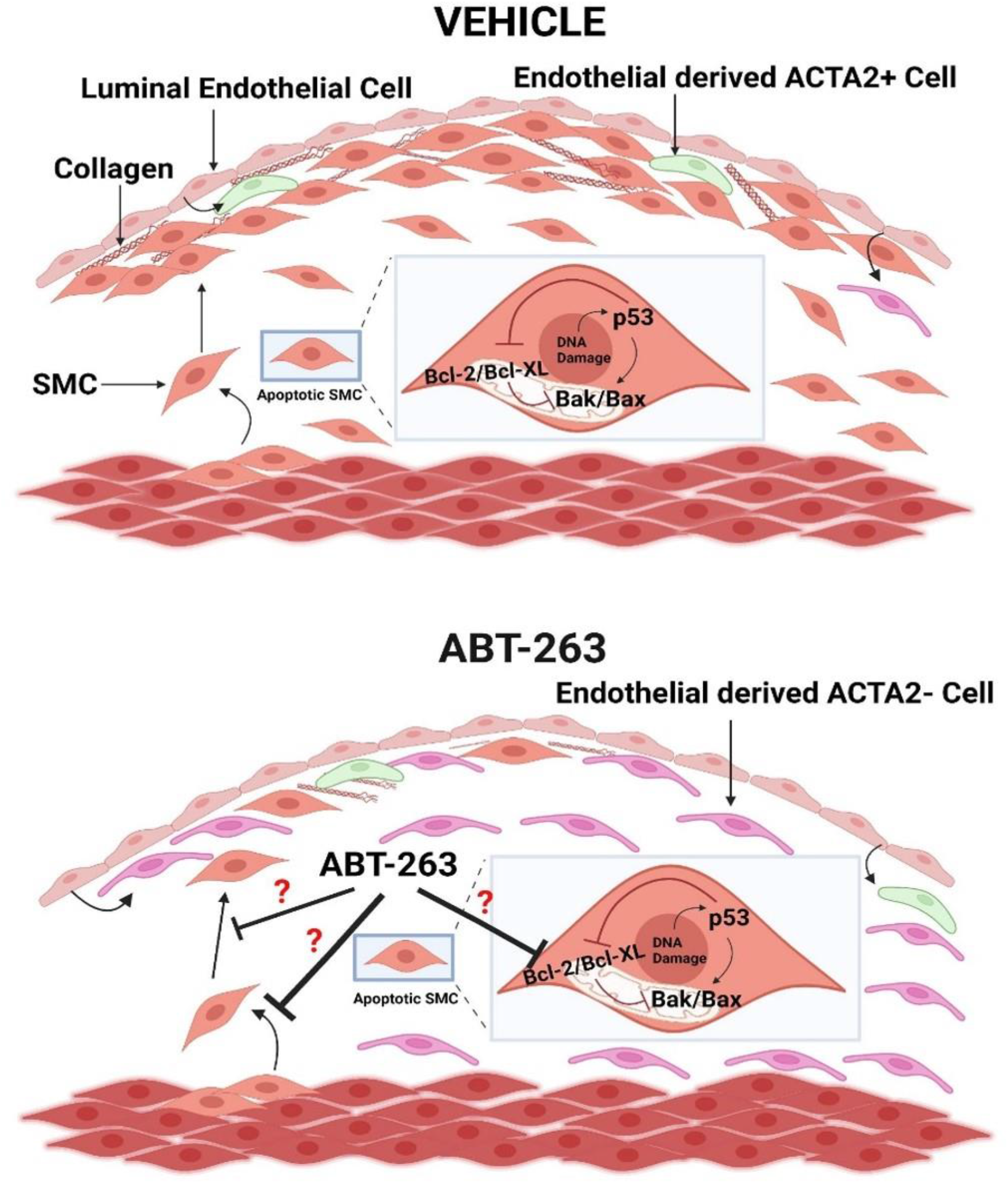

**Highlights:**

- Treatment of Apoe-/- mice with advanced atherosclerosis with the senolytic agent ABT-263 increased mortality by >50%.
- ABT-263 showed a 90% reduction in SMC but a 60% increase in endothelial cell (EC) contributions to lesions via EC to mesenchymal transition (EndoMT) but prevented adaptive increases in investment of EC-derived cells into the fibrous cap via beneficial EndoMT to myofibroblast transitions that we have shown normally occur when SMC investment into fibrous cap of lesions is impaired.
- Knock out (KO) of Klf4 in SMC, which results in smaller but more stable atherosclerotic lesions, was associated with reduced expression of pro-senescence markers, but preserved expression of the anti-senescence marker, telomerase reverse transcriptase although it is unclear if the latter is causal or an effect.

## Introduction

The rupture or erosion of advanced atherosclerotic plaques resulting in ischemic heart disease or stroke are the leading causes of death worldwide^1^. Human histopathological studies have clearly established that lesions containing a thick extracellular matrix (ECM)-rich fibrous cap and an abundance of α-SMA^+^ versus CD68^+^ cells are less likely to rupture^2^. Most investigators in the field have assumed that α-SMA^+^ lesion cells are derived from SMC and have a beneficial role in lesion pathogenesis by being the exclusive source of ECM-producing fibrous cap cells^3^. In addition, they assumed that CD68+ cells are macrophages that exacerbate lesion pathogenesis by increasing inflammation and giving rise to foam cells. However, subsequent studies by our lab^4^ and many others^5–8^ have either refuted several key aspects of these assumptions or shown that they are overly simplistic. This includes the following. **First**, α-SMA^+^ staining is highly unreliable for identifying SMC-derived cells in lesions given that nearly 80% of SMC-derived lesion cells lack detectable expression of this marker and activate markers of other cell types including macrophages^4^. In addition, we recently showed that 40% of α-SMA^+^ fibrous cap cells are derived from a non-SMC including EC that have undergone EC to mesenchymal transition (EndoMT) and macrophages undergoing macrophage to myofibroblast transition (MMT) in advanced BCA lesions from *Apoe*^-/-^ and *Ldlr* deficient mice fed a WD for 18 weeks^9^. **Second**, there are qualitative differences in plaque stability depending on the source of α-SMA^+^ fibrous cap cells with increased EndoMT and MMT only being able to transiently compensate for the loss of SMC investment into lesions induced by SMC knockout (KO) of *Pdgfβr* or lethal irradiation^9^. **Third,** there is compelling evidence showing that each of the major cell types involved in atherosclerosis including macrophages^10^, T-Cells^11^, B-cells^12^, neutrophils^13^, endothelial cells^9^, and SMC^4, 14–16^ can have beneficial or detrimental effects on lesion development or late-stage pathogenesis depending on the nature of their phenotypic transitions^17^. For example, using a combination of SMC lineage tracing, SMC-specific KO of *Klf4* or *Oct4*, and scRNAseq analysis of lesions, we showed that SMC can have a beneficial or detrimental role in lesion pathogenesis^15^. Results with SMC KO of *Klf4* were particularly intriguing since this resulted in what would be the ideal therapeutic outcome in patients at risk for development of atherosclerosis. Specifically, SMC-*Klf4* KO lesions were 50% smaller and exhibited features associated with increased plaque stability including a two-fold increase in the thickness of the extracellular matrix (ECM) rich α- SMA^+^ fibrous cap^4^. *Klf4* was subsequently identified as a coronary artery disease GWAS variant^18^. Moreover, our lab recently showed that loss of *Klf4* in SMCs resulted in a marked reduction of mRNA transcripts associated with cellular senescence and inflammation including >40 putative *KLF4* target genes previously identified as coronary artery disease GWAS variants. Taken together, results suggest that the beneficial effects of SMC *Klf4* KO in mouse models, and for *Klf4* being a CAD GWAS variant are due in part to its role in enhancing cell senescence and inflammation.

Senescence is a term used to define a state of irreversible growth arrest and includes replicative senescence, or stress-induced premature senescence (SIPS)^19^. Indeed, previous studies have shown that the atherosclerotic lesions in humans and mice contain relatively large numbers of senescent cells^20–22^, and claim that these senescent cells promote atherosclerosis and contribute to destabilization of plaques^23^. However, the origins and functional properties of these senescent cells are uncertain since they relied on methods that eliminated “senescent cells” based on expression of a single marker gene which is not specific for senescent cells, and/or relied on one or two traditional, albeit unreliable, markers to identify SMC-, EC- and macrophage-derived cells within lesions^23, 24^. For example, when Childs et al.^25^ reduced senescent cells using a ganciclovir (GCV) dependent-*p16* driven thymidine kinase genetic approach to induce death of p16^Ink4a+^ cells they discovered reduced Sudan-IV^+^ plaque burden and increased fibrous cap thickness of atherosclerotic lesions within the descending aorta of *Ldlr*^-/-^ female mice. This same study showed that treatment of *Ldlr*^-/-^ female mice at a dosage of 100 milligram/kilogram/body weight (mg/kg/bw) of the senolytic drug ABT-263 *(Navitoclax)* throughout 88 days of high-fat diet feeding reduced the Sudan-IV^+^ area of the aorta^25^. In addition, a more recent study by Childs et al.^26^ showed that aortic arch lesions of Myh11 CreER^T2^ TdTomato *Ldlr*^-/-^ male mice fed a WD for 14 weeks followed by 9 weeks of a low fat diet and treatment with ABT-263 had a reduced proportion of tdTomato^+^ cells that expressed the osteochondrogenic marker *Runx1* as compared to vehicle treated mice^26^. ABT-263 inhibits interactions between the anti-apoptotic protein BCL-X^L^ and the pro apoptotic protein BAX thus enhancing clearance of senescent cells by reducing their capacity to evade efferocytotic clearance^27^. ABT-263 disrupts *Bcl-2/Bcl-xL* interactions with pro-death proteins (e.g., Bim), leading to the initiation of apoptosis^39^. Thus, Childs et al.,^25, 26^ interpreted their results as evidence that the beneficial effects on atherosclerosis observed with ABT-263 treatment were the direct result of enhanced clearance of senescent cells from lesions. In recent years these results have been challenged by results of studies by Martin Bennett and co-workers who showed the following^28^. **First**, they found that “presumed” selective clearance of senescent cells using a p16 driven thymidine kinase genetic approach similar to Childs et al.^25^ did not reduce Oil red O^+^ plaque burden, aortic root lesion size, fibrous cap thickness, or necrotic core area in *Apoe*^-/-^ mice (males and females combined), but instead increased apoptotic cells and induced inflammation. **Second**, treatment of *Apoe*^-/-^ male and female mice with established lesions with 50 mg/kg/bw of ABT-263 did reduce aortic root lesion size but did not increase cap thickness. **Third**, ABT-263 treatment of *Apoe*^-/-^ mice did not reduce relative mRNA expression of *p16* and senescence-associated secretory phenotypes, including *IL-6, IL-1α, Tnf-α, IL-18* and *Mmp-12* in aortic arch tissue, but increased thrombocytopenia and reduced monocyte and leukocyte populations. Another independent study from James Kirkland’s lab showed that treatment of *Apoe*^-/-^ mice with the senolytic agent Dasatinib *plus* Quercetin did not reduce lesion size, but reduced plaque calcification^29^. The reasons for the disparate results of the preceding studies are unclear but may be related to male versus female sex, *Apoe*^-/-^ versus *Ldlr*^-/-^ mice, and differences in the senolytic drugs tested and/or dosing regimens. However, a common limitation of these previous studies was the failure to rigorously define the effects of ABT 263 on SMC despite each study claiming this is the major source of senescent cells. In addition, there have been no studies of the effects of ABT-263 on endothelial cells (EC) which along with SMC comprise 90% of α-SMA^+^ myofibroblast-like cells in the protective fibrous cap. These studies also include minimal assessment of indices of plaque stability and were largely focused on assessment of aortic root lesions, which poorly replicate the histology of human lesions. As such, there are still major uncertainties regarding the effects of senolytic therapies on the cellular composition of advanced atherosclerotic lesions, the phenotypes of the lesion cells, and overall indices of plaque stability that need to be clarified before determining their usefulness in treating patients with advanced atherosclerotic disease.

*Studies herein test two hypotheses. First, we hypothesized that the beneficial effects of SMC KO of Klf4 are due in part to reduced cell senescence and an associated reduction in inflammation. Second, we hypothesized that treatment of Western Diet (WD) fed SMC- and EC lineage tracing Apoe*^-/-^ *mice with the senolytic agent ABT-263 will result in loss of senescent cells from atherosclerotic lesions thereby reducing lesion inflammation and size, as well as inducing changes consistent with increased plaque stability including a thicker SMC- and ECM-rich* α-SMA+ *protective fibrous cap.* Consistent with our first hypothesis, KO of *Klf4* in SMC in WD fed *Apoe*^-/-^ mice was associated with marked reductions in plaque size, lipid deposition, and the overall prevalence of senescent cells. However, contrary to expectations, the results of studies testing our second hypothesis showed that treatment of WD-fed *Apoe*^-/-^ mice with ABT 263 had multiple detrimental effects including inducing marked reductions in Myh11-eYFP^+^ SMC-derived cells within lesions and the fibrous cap, reduced α-SMA^+^ fibrous cap thickness, increased EndoMT, and increased mortality.

## Methods

### Mice

The University of Virginia Animal Care and Use Committee approved animal protocols (Protocol 2400). The Myh11-CreER^T2^-eYFP and Cdh5-CreER^T2^-eYFP *Apoe*^-/-^ mice used in the intervention study have been described in our previous studies^4,9^. In addition, Myh11-CreER^T2^-eYFP *Klf4* WT vs KO *Apoe*^-/-^ mice were used for the studies related to loss of *Klf4* in SMC experiments as previously reported^4^.

#### Diet and treatment: ABT-263 intervention studies on advanced atherosclerotic mice

In Myh11-CreER^T2^-eYFP and Cdh5-CreER^T2^-eYFP *Apoe*^-/-^ mice, Cre recombinase was activated with a series of ten tamoxifen injections (1mg/day/mouse; Sigma Aldrich, T-5648) over a 2-week period. One week after the tamoxifen treatment, mice were switched from a normal chow diet (Harlan Teklad TD.7012) to a high fat Western-type diet (WD), containing 21% milk fat and 0.15% cholesterol (Harlan Teklad; TD.88137) for 18 weeks followed by 100mg/kg/bw ABT-263 treatment for 6 (2 cycles 5 days ON and 14 days OFF) weeks or 50mg/kg/bw ABT-263 treatment for 9 (3 cycles 5 days ON and 14 days OFF) weeks. ABT-263 (S1001) was obtained from Selleck (Houston, TX 77014, USA), formulated in Vehicle (PBS with 15% dimethylsulfoxide and 7% Tween-20) and injected intraperitoneally (IP) at a dose of 100mg/kg/bw or 50mg/kg/bw as shown in the figures, figure legends, and results sections.

### Atherosclerotic plaque morphometry

Paraformaldehyde-fixed paraffin-embedded BCAs were serially cut into 10 μm thick sections from the aortic arch to the bifurcation of the right subclavian artery. For morphometric analysis, we performed modified Russell-Movat staining on three locations along the BCA at 150 μm, 450 μm, and 750 μm, from the aortic arch as previously described^9^. The lesion, lumen, external elastic lamina (outward remodeling), and necrotic core areas as well as the internal elastic lamina area were measured on digitized images of the Movat staining using Fiji (Image J) software. Picrosirius Red staining was performed to assess collagen content and digitized images of the picrosirius staining was measured using Fiji software. Masson trichrome staining was performed to assess the fibrous tissue in the liver sections.

### Immunofluorescent staining

BCA sections were de-paraffinized and rehydrated in xylene and ethanol series. After antigen retrieval (H-3300-250, Vector Laboratories, Newark, CA 94560, USA), sections were blocked with fish skin gelatin–PBS (6 g/L) containing 10% horse serum for 1 hour at room temperature. Slides were incubated with the following antibodies: mouse monoclonal smooth muscle α-actin-Cy3 (α-SMA) (4.4 μg/mL, clone 1A4, C6198, Sigma Aldrich, Rockville, MD 20850, USA), goat polyclonal anti-GFP (4 μg/mL, ab6673, Abcam, Waltham, MA 02453, USA) for detection of eYFP, rabbit polyclonal *p21* (18 μg/mL, 10355-1-AP, Proteintech, Rosemont, IL 60018, USA), TUNEL (Cat 30074 Bioutium, Fremont, CA, USA). The secondary antibodies donkey anti-goat conjugated to Alexa 488 (5 μg/mL), donkey anti-rabbit conjugated to Alexa 647 (A31573 5 μg/mL), DAPI (0.05 mg/mL, D3571,) was used as a nuclear counterstain and slides were mounted using Prolong Gold Antifade (P36930) were purchased from Thermo Fisher Scientific, Waltham, MA 02451, USA.

### Imaging

Movat and picrosirius red staining of brachiocephalic arteries and masson trichrome staining of liver sections were imaged by using a Leica thunder imager microscope. Image acquisition was performed with Leica software. Digitized images were analyzed with Fijisoftware. Immunofluorescent staining was imaged using a Zeiss LSM880 airy scan confocal microscope to acquire a series of z-stack images at 1-μm intervals. Zen 2009 Light Edition Software (Zeiss) was used for the analysis of each z-stack image and single cell counting was performed for phenotyping and quantifying the cell population comprised within the 30μm thick layer proximal to the lumen (i.e., fibrous cap area). Assessment of α-SMA^+^ cap thickness (normalized to lesion) was performed using Zen 2009 Light Edition Software. Maximal intensity projection of representative images wereused to generate the representative images included in the figures.

### Western blotting

The method has been described previously^30^. The primary antibodies used for p16 were from Abcam, 2μg/mL, Ab189034, Waltham, MA 02453, USA), Klf4 (1 μg/mL, CST4038T) and β-actin (1 μg/mL, 4967S) from Cell Signaling Technology, Danvers, MA 01923, USA.The anti-Rabbit IgG (A4914) HRP-conjugated secondary antibody was purchased from Sigma Aldrich, Rockville, MD 20850, USA.

### Senescence-Associated β–Galactosidase (SAβG) Staining

SaβG staining of aortas for senescent cells was examined using a Senescence detection assay kit (Millipore-Sigma Cat No.KAA002, Rockville, MD 20850, USA)^30^. Freshly isolated aortas were kept on ice in a 12 well plate containing PBS until all the aortas were harvested. Aortas were washed twice with the PBS, and then fixed with the diluted fixation solution provided in the kit for 10 minutes as per the manufacturer’s instructions. After the fixation, aortas were washed thrice with PBS and then 1mL of freshly prepared SAβG solution was added to a well of 12 well plates containing one aorta per well. Then, plates were wrapped with aluminum foil to avoid light exposure and placed in a 37^0^C incubator for 24h. After that, aortas were washed thrice with PBS, and photographs were taken with a mobile phone camera.

### Statistics

Statistics were performed using GraphPad prism 9. Data are presented as mean ± SEM. Animal numbers and type of statistical analysis done are reflected within figures and figure legends. A p-value ≤ 0.05 was considered statistically significant.

## Results

### *Klf4* knockout in SMC resulted in reduced overall lesion senescence

As an initial test of our hypothesis *that the multiple beneficial effects of SMC KO of Klf4 are due in part to reduced cell senescence,* male SMC *Klf4* WT and KO *Apoe*^-/-^ mice were fed a WD for 18 (Figure I in the data supplement) or 26 weeks (Fig.1A) and aortas were then stained for senescence-associated-β-gal (SAβG). Interestingly, SMC *Klf4* WT mice aortas showed highly prevalent SAβG^+^ senescent cells in the brachiocephalic artery (BCA), aortic root, aortic arch, and abdominal aorta (Fig.1B). In contrast, there was almost no SAβG^+^ staining in SMC *Klf4* KO mice (Fig.1B) as seen in whole aortic preparations and in cross sections of the thoracic and abdominal aorta counter stained with nuclear fast red (Fig.1Bi-iv). Similar results were found in mice fed WD for 18 weeks (Figure IB-IC in the data supplement). Loss of *Klf4* in SMC was also associated with an increase in the Telomere Reverse Transcriptase (*Tert*^+^) anti senescence marker, but a decrease in *p16*^+^ (pro-senescence marker) clusters, as assessed from uniform manifold approximation and projections (UMAPs) of scRNAseq data sets of BCA lesions from SMC *Klf4* WT and KO mice^15^ (Figure ID-IE in the data supplement). KO of *Klf4* in SMC preserved the *Tert*^+^ cells in most of the clusters of lesion cells as shown by UMAP analysis (Figure ID in the data supplement). In contrast, KO of *Klf4* in SMC reduced the number of SMC-derived cells positive for the pro-senescence marker *p16* (*Cdkn2a*) (Figure IE in the data supplement). Indeed, our previous study showed that *Klf4* binds to Telomeric RNA (*TERC*) in SMCs of advanced atherosclerotic lesions based on chromatin immunoprecipitation (CHIP) assays^4^. Next, cultured murine aortic SMC were transfected with siKlf4 to determine if knock down (KD) of *Klf4* in SMC reduces the pro-senescent marker *p16* directly or through crosstalk with other cell types in the lesion. Results showed that KD of *Klf4* reduced *p16* expression in SMC at both the mRNA and protein levels (Figure IF-IG in the data supplement).

**Figure 1.**
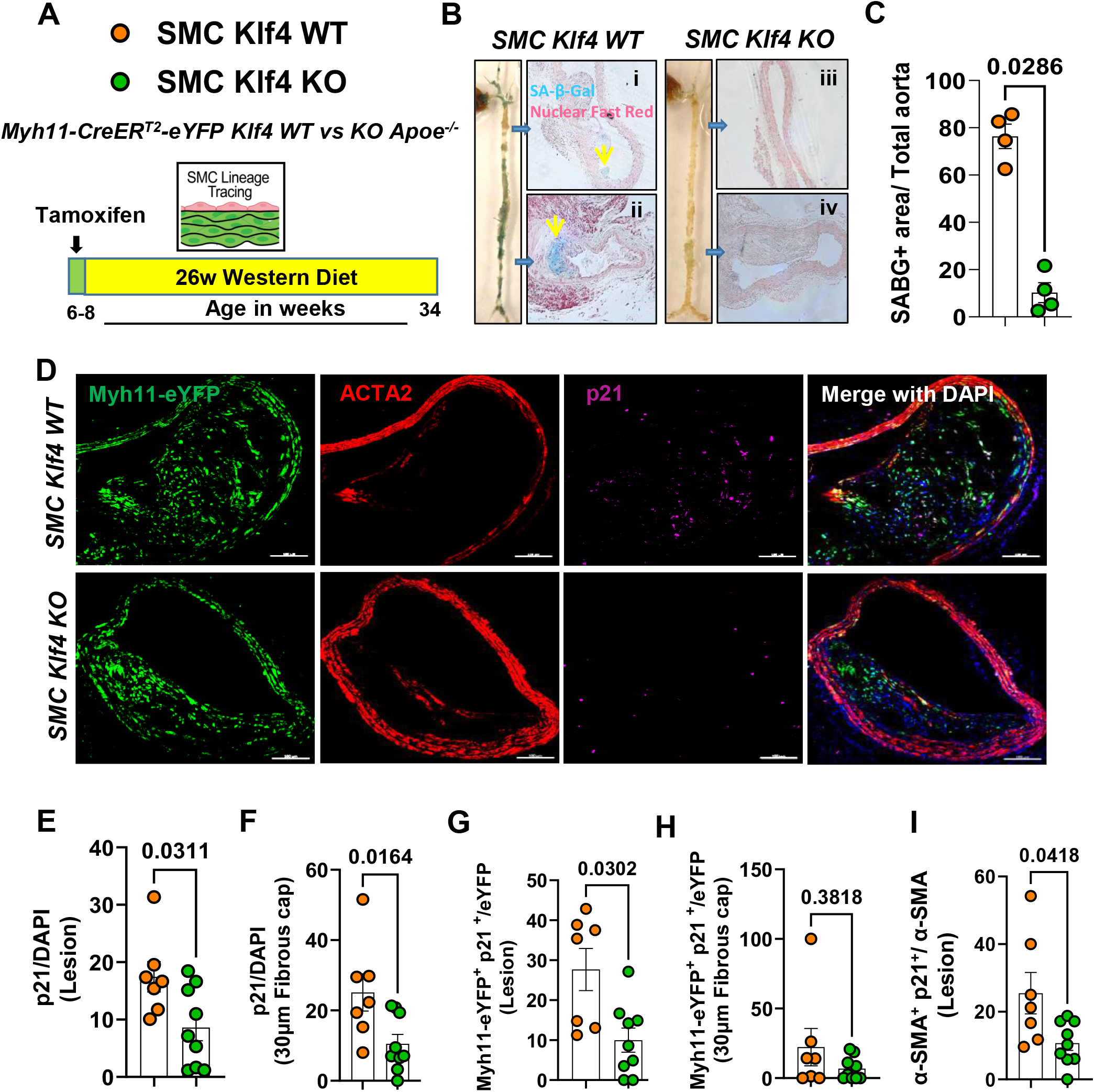
SMC-Klf4 KO resulted in a marked reduction in overall lesion senescence including reduced expression of the pro-senescence markers *p21*, *p16*, and SAβG, but preserved anti-senescence marker telomerase reverse transcriptase positive cells. (A) Experimental design, SMC ((Myh11-CreER^T2^-eYFP)) lineage tracing *Klf4* WT vs KO *Apoe*^-/-^ mice were injected with tamoxifen at 6 to 8 weeks of age and subsequently placed on Western diet (WD) for 26 weeks to induce advanced atherosclerosis. **(B)** SAβG staining was performed as described in the methods section on freshly isolated aortas. (**Bi-iv**) thoracic and abdominal aortic sections from SAβG stained aortas from *B* and counter stained with nuclear fast red. **(C)** Quantification of SAβG+ area of the aorta. **(D)** Representative images of co-staining for eYFP (for detecting SMC), α-SMA, *p21* (a marker of senescence) and DAPI (nucleus) in advanced BCA lesions from SMC *Klf4* WT and KO animals fed a WD for 26 weeks. The confocal images show a maximum intensity projection ×20 zoom with a scale bar of 100µm. Quantification of the frequency of *p21*^+^ (p21^+^/DAPI) senescent cells in the lesion **(E)** and fibrous cap **(F)** as a percent of total cells in the lesions. Quantification of SMC derived *p21*^+^ (eYFP^+^ p21^+^/eYFP) senescent cells in the lesion **(G)** and the fibrous cap **(H)** as a percent of total SMC. Quantification of α-SMA^+^ senescent (α-SMA^+^ p21^+^/ α-SMA) cells in the lesion **(I)** as a percentage of total α-SMA^+^ cells. Mann-Whitney U-tests were used in C, and E-I. The error bars show the standard error of the mean (SEM). Independent animals are indicated as individual dots (WT, n=7, and KO, n=9). The p-values are indicated on the respective graphs.

A previous study from our lab showed that *Klf4* is a suppressor of SMC differentiation markers and an inhibitor of SMC proliferation by inducing *p21*^WAF1/Cip1^ expression in concert with *p53*^31^. Both *p53* and *p21* are widely accepted pro-senescence markers^32^. Therefore, we stained BCA sections of SMC *Klf4* WT and KO mice fed a WD for 18 weeks with a *p21* antibody to determine if SMC *Klf4* KO would reduce the frequency of p21^+^ senescent cells in atherosclerotic lesions (Fig.1D). As predicted, there was a reduction in *p21*^+^ senescent cells (p21/DAPI) in the BCA lesions and fibrous cap of SMC *Klf4* KO versus WT control mice (Fig.1E and 1F). The fraction of SMC-derived *p21*^+^ cells (Myh11-eYFP^+^ p21^+^/eYFP) was also reduced in the lesions but not in the fibrous cap of these same mice (Fig.1G-1H). Furthermore, the fraction of α-SMA^+^ cells which expressed the senescence marker *p21* was also reduced in the lesions of SMC *Klf4* KO versus WT control mice (Fig.1I). Results show that KO of *Klf4* in SMC was associated with a marked reduction in the prevalence of senescent cells within the aorta and BCA lesions of *Apoe*^-/-^ mice fed WD for 18 or 26 weeks.

Taken together, the preceding results extend results of previous studies from our lab^4^,^15^ showing that SMC *Klf4* KO has multiple beneficial effects including resulting in smaller lesions with a thicker α-SMA^+^ fibrous cap^4^ containing reduced numbers of senescent cells (Figure 1 and supplemental figure I from present study). However, it is not clear if the reduction in senescent cells contributes causally to the beneficial effects on lesions or if the reduced number of senescent cells is secondary to loss of *Klf4* in SMC acting through other mechanisms. To distinguish these possibilities, we sought alternative approaches to reduce the prevalence of senescent cells within advanced lesions.

### Treatment of *Apoe*^-/-^ mice with advanced atherosclerotic lesions with the senolytic drug ABT-263 reduced the number of SMC within BCA lesions and was associated with increased EndoMT derived lesion cells and mortality

To determine if increased clearance of senescent cells could induce beneficial changes in late stage complex atherosclerotic lesions, SMC- and EC-lineage tracing *Apoe*^-/-^ mice were fed a WD for 18 weeks and then treated with 3 cycles (Fig.2A) of 100mg/kg/bw ABT-263 as used previously by Childs et al.^25^. We elected to do an intervention rather than a prevention study to better model clinical scenarios in patients with advanced disease^33^. However, the experiment had to be unexpectantly terminated for ethical reasons, as per our ACUC approved animal protocol, after two cycles of ABT 263 because >50% of the ABT-263 treated mice died (Fig.2B). ABT-263 treated mice had no changes in BCA lesion size (Fig.2C-2D), lumen area and outward remodeling (EEL external elastic lamina area) (Fig.2E-2F), and no change in aortic root lesion area (Fig.2G-H). However, ABT-263 treated mice showed multiple detrimental changes in lesion composition including reduced BCA lesion collagen content (Fig.2I-2J), a reduced α-SMA^+^ fibrous cap area (Fig.3A-3C), and a reduced fraction of α-SMA^+^ or α-SMA^-^ SMC derived cells (Myh11-eYFP^+^/DAPI) in lesions and the 30um fibrous cap area (Fig.3B and 3D-3E, and Figure IIA-IIC in the data supplement). Surprisingly, ABT-263 treatment did not change the frequency of apoptotic cells (TUNEL^+^/DAPI) in the BCA lesions as would be expected based on its mechanism of action (Figure IIIA-C in the data supplement). To determine if the non-SMC-derived α-SMA^+^ fibrous cap cells in this study were derived from EC, we treated EC-lineage tracing mice with the same dose of ABT-263 and assessed (Fig.3F) BCA lesions for eYFP, α-SMA and DAPI (Fig.3G). The ABT-263 treatment also reduced the α-SMA^+^ cap area and the fraction of α-SMA^+^ (α-SMA^+^/DAPI) cells in the 30um fibrous cap area of these mice (Fig.3H and Figure IID-IIE in the data supplement). Interestingly, ABT-263 treatment increased endothelial cells (Cdh5-eYFP^+^/DAPI) in the fibrous cap and lesions but did not increase endothelial-derived α- SMA^+^ (Cdh5-eYFP^+^ α-SMA^+^/ α-SMA^+^) cells (Fig.3I-3J and Figure IIF-IIH in the data supplement). Taken together, results show that ABT-263 treatment at 100mg/kg/bw had no effect on lesion size but markedly reduced the probability of survival. This treatment regimen was also associated with major detrimental changes in lesion composition including reduced α-SMA^+^ fibrous cap thickness and SMC investment into the fibrous cap. In addition, ABT-263 treatment prevented adaptive increases in investment of EC-derived cells into the fibrous cap via EndoMT to myofibroblast transitions that we have shown normally occur when SMC investment into the fibrous cap of lesions is impaired^9^.

**Figure 2.**
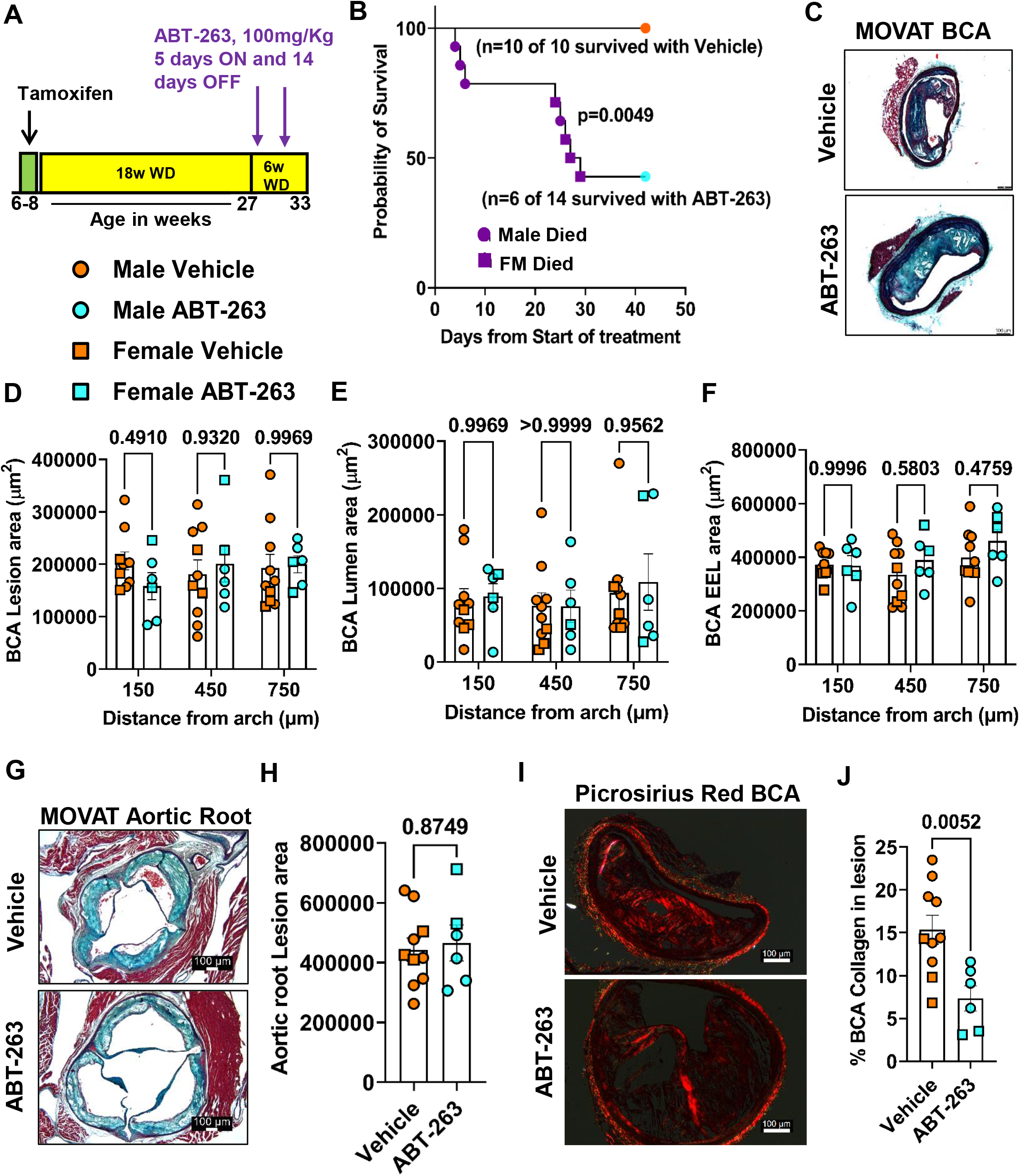
Treatment of SMC (Myh11-CreER^T2^-eYFP) and EC (Cdh5-CreER^T2^-eYFP)-lineage tracing Apoe^-/-^ mice with advanced atherosclerotic lesions with ABT-263 had no effect on lesion size but increased mortality. **(A)** Experimental design, SMC- and EC-lineage tracing *Apoe*^-/-^ mice were fed a WD for 18 weeks followed by ABT-263 treatment on western diet (WD) for 6 weeks. **(B)** Probability of survival (Kaplan-Meier curve). **(C)** Representative 10x images with 100μm scale bar of MOVAT staining on Brachiocephalic Artery (BCA). **(D)** Lesion area from D. **(E)** Lumen area from D. **(F)** External elastic lamina (EEL) area from D, for outward remodeling. **(G)** Aortic roots stained with MOVAT. **(H)** Lesion area quantification from G. **(I)** Representative 10x images with 100μm scale bar of Picrosirius red staining on Brachiocephalic Artery (BCA). **(J)** Quantification of matured (red) collagen content normalized to lesion area from figure I. A repeated measures Two-way ANOVA method was used for statistical analysis in D-F whereas Mann-Whitney U-tests were used for H and J. Biologically independent animals are indicated as individual dots, error bars show the SEM, Mantel-Cox test used for B. The p-values are indicated on the respective graphs.

**Figure 3.**
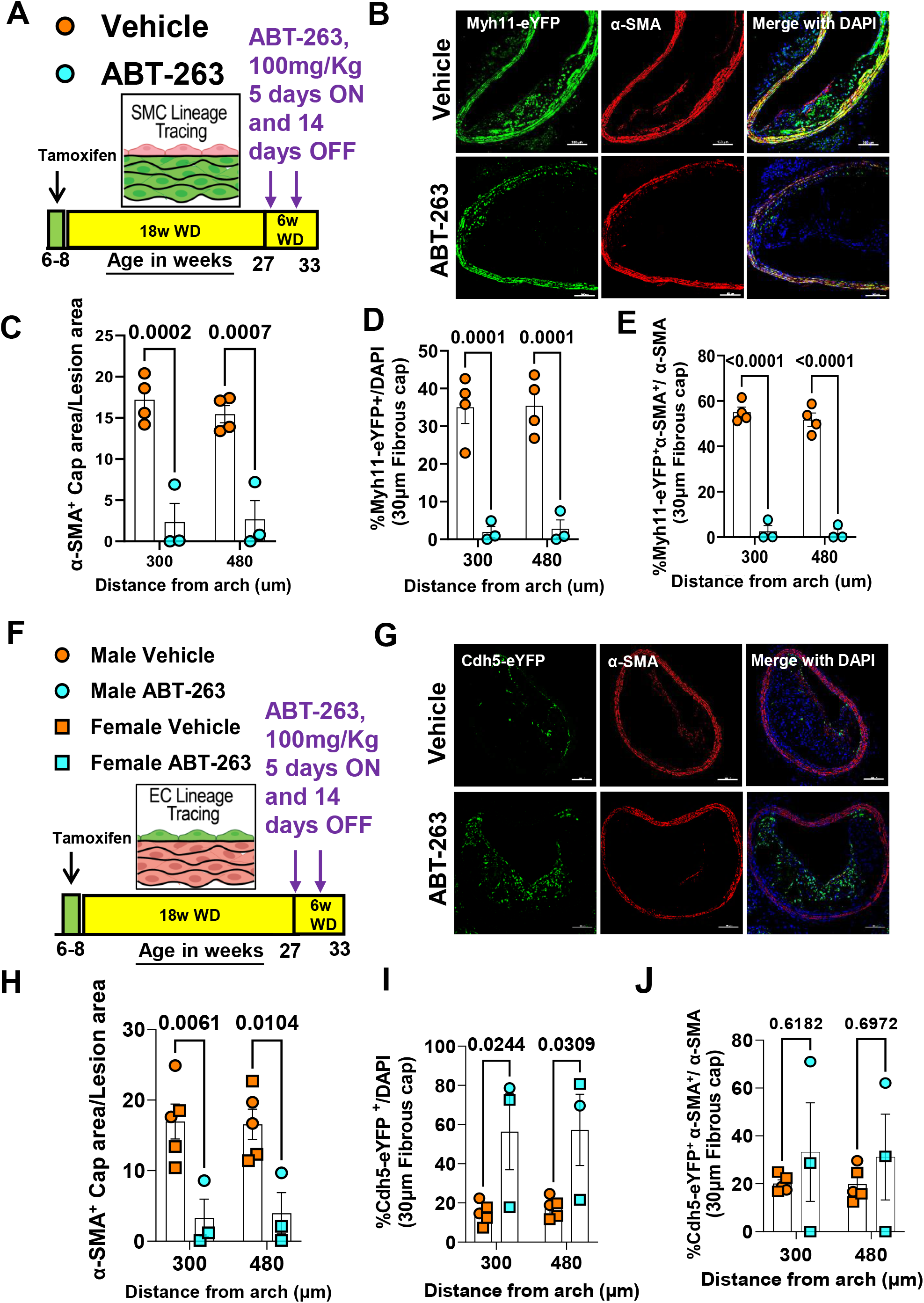
Treatment of SMC (Myh11-CreER^T2^-eYFP) and EC (Cdh5-CreER^T2^-eYFP) lineage tracing *Apoe*^-/-^ mice with advanced atherosclerotic lesions with the senolytic drug ABT-263 (100mg/kg/bw) was associated with a marked reduction in the number of SMC within BCA lesions but an increase in EC-derived cells undergoing EndoMT although the latter did not result in increased investment of EC-derived α-SMA+ cells into the fibrous cap. **(A)** Experimental design for figures *B-E*, SMC-lineage tracing *Apoe*^-/-^mice were fed a WD for 18 weeks followed by 100mg/kg/bw ABT-263 treatment on WD for 6 weeks. **(B)** Representative confocal images of co-staining for eYFP (for detecting SMC), α-SMA^+^, and DAPI in advanced BCA lesions from experiment *A*. The confocal images show a maximum intensity projection ×20 zoom and a scale bar of 100µm. **(C)** α-SMA^+^ cap area normalized to lesion area (α- SMA^+^ cap area/Lesion area). **(D)** Quantification of the percentage of SMC-derived (Myh11-eYFP^+^/DAPI^+^) cells in the fibrous cap, and **(E)** quantification of the % SMC derived α-SMA^+^ (Myh11-eYFP^+^ α-SMA^+^/α-SMA^+^) cells in the fibrous cap. **(F)** The experimental design for figures *H-J*, EC-lineage tracing *Apoe*^-/-^ mice were fed a WD for 18 weeks followed by 100mg/kg/bw ABT-263 treatment on WD for 6 weeks. **(G)** Probability of survival (Kaplan-Meier curve). **(H)** Representative confocal images of co-staining for eYFP (for detecting EC), α-SMA^+^, and DAPI in advanced BCA lesions from experiment *F*. **(I)** Quantification of the percentage of EC-derived Cdh5-eYFP^+^ DAPI^+^ cells in the fibrous cap, and **(J)** quantification of the percentage of EC-derived α-SMA^+^ (Cdh5-eYFP+ α-SMA^+^/ α-SMA^+^) cells in the fibrous cap. The two-way ANOVA method was used for statistical analyses in C-E and H-J. Biologically independent animals are indicated as individual dots. Error bars are shown with the SEM. A Mantel-Cox test was used for G. The p-values are indicated on the figures.

### *Apoe*^-/-^ mice with advanced lesions treated with a reduced dose of ABT-263 also had increased mortality and detrimental changes in lesion composition

Given the multiple detrimental effects of treating WD-fed *Apoe*^-/-^ mice with ABT 263 at 100mg/kg/bw we repeated the preceding studies with 50mg/kg/bw of ABT-263. EC-lineage tracing *Apoe*^-/-^ mice were fed a WD for 18 weeks followed by treatment with ABT-263 at half our original dose (i.e., 3 cycles of 50mg/kg/bw) (Fig.4A). Contrary to our expectations, even this lower dose of ABT-263 was associated with increased mortality (Fig.4B). Similar to the higher dose of ABT-263, there were no changes in lesion size, lumen area, outward remodeling, and necrotic core area of BCA lesions (Fig.4B-G). There were also no reductions in aortic root lesion area (Figure IIID-IIIF in the data supplement) in male or female mice. Surprisingly, the lower dose of ABT-263 reduced collagen content in male *Apoe*^-/-^ mice (Fig.4H-4I). Unlike the 100mg/kg dose, treatment at the lower dose did not reduce α-SMA^+^ cap thickness, area, or α-SMA^+^ cells in the fibrous cap or lesions (Fig. 4J-4L and Figure IVA-IVB in the data supplement). The lower dose of ABT-263 was associated with increased EndoMT (Cdh5-eYFP^+^/DAPI) in male *Apoe*^-/-^ mice but no change in endothelial-derived α-SMA^+^ (Cdh5-eYFP^+^ α-SMA^+^/ α-SMA) cells in the lesions or 30um fibrous cap area (Fig.4M-4N and Figure IVC-IVD in the data supplement). Taken together, even the reduced dose of ABT-263 increased mortality and did not show beneficial changes in plaque size and composition.

**Figure 4.**
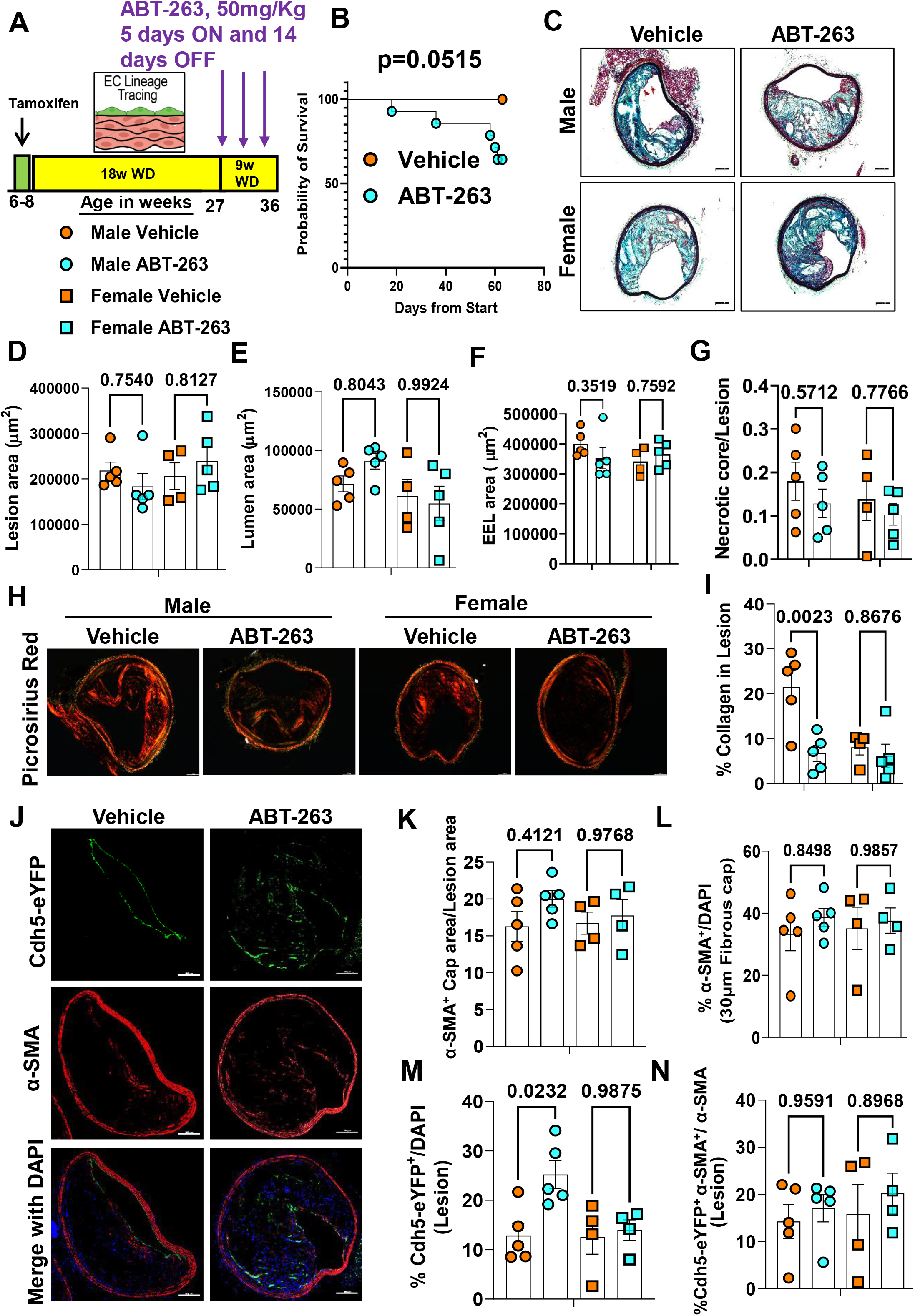
263 Treatment of Apoe^-/-^ mice with advanced lesions with a reduced dose (50mg/kg/bw) of ABT decreased collagen content within BCA lesions and was also associated with increased mortality. **(A)** Experimental design, EC-lineage tracing *Apoe*^-/-^ mice were fed a WD for 18 weeks followed by 50mg/kg/bw ABT-263 treatment on WD for 9 weeks. **(B)** Probability of survival (Kaplan-Meier curve). **(C)** Representative 10x images with 100μm scale bar of MOVAT staining of the BCA. **(D)** Lesion area from C. **(E)** Lumen area from C. **(F)** External elastic lamina (EEL) area from C, for outward remodeling. **(G)** Necrotic core area normalized to lesion size. **(H)** Representative 10x images with 100μm scale bar of Picrosirius red staining on Brachiocephalic Artery (BCA). **(I)** Quantification of mature (red) collagen content normalized to lesion area from figure H. **(J)** Representative confocal images of co-staining for eYFP (for detecting EC), α- SMA^+^, and DAPI in advanced BCA lesions. **(K)** α-SMA^+^ cap area normalized to lesion area (α-SMA^+^ cap area/Lesion area). **(L)** Quantification of the percentage α-SMA^+^ (α- SMA^+^/DAPI) cells in the fibrous cap. **(M)** Quantification of the percentage EC-derived (Cdh5-eYFP^+^/DAPI^+^ cells in the lesion, and **(N)** quantification of the percentage EC derived α-SMA^+^ (Cdh5-eYFP^+^ α-SMA^+^/α-SMA^+^) cells in lesions. The two-way ANOVA method was used for statistical analysis in D, G, and K-L, and biologically independent animals are indicated as individual dots. Error bars show the SEM. A Mantel-Cox test used for statistical analysis in B. The p-values are indicated on the respective graphs.

### Low dose ABT-263 treatment of WD-fed *Apoe*^-/-^ mice reduced plasma *Cxcl5* and was associated with an abnormal fibrous liver phenotype

To determine if ABT-263 treatment reduces senescence-associated secretory phenotypes (SASPs) and pro-inflammatory cytokine levels, we performed luminex assays on plasma samples. Here we discovered that ABT-263 did not reduce SASPs and cytokines including, *IL-1β, IL-1α, IL-6, Mcp-1, Tnf-α, and Ifn-γ (*Figure V in the data supplement). However, ABT-263 treatment reduced *LIX* (*Cxcl5*) levels in female *Apoe*^-/-^ mice (Figure V in the data supplement). *Cxcl5* has an atheroprotective role in that inhibition of *Cxcl5* has previously been shown to induce significant macrophage foam cell accumulation in murine atherosclerotic plaques^34^.

Mice treated with ABT-263 also showed an abnormal liver phenotype with increased Masson trichrome positive fibrous tissue. However, ABT-263 treatment did not result in changes in alanine aminotransferase (ALT) and aspartate aminotransferase (AST) levels in the plasma (Figure 5F-5G) suggesting that the ABT-263 induced increase in liver damage and fibrosis may be associated with other factors or mechanisms. Although ABT 263 did not reduce lesion size, it reduced cholesterol and LDL levels in male but not female mice *Apoe*^-/-^ mice (Figure VIB-VIC in the data supplement). Triglycerides levels were not changed with ABT-263 treatment (Figure VID in the data supplement). ABT-263 treatment did not increase thrombocytopenia (Figure VIE in the data supplement). Moreover, ABT-263 did not result in changes in basophils, lymphocytes, WBC, RBC, nucleated RBC, hemoglobin, monocytes, and neutrophils in the blood (Figure VII in the data supplement). Also, ABT-263 treatment did not change the body weights (Figure VIII in the data supplement).

**Figure 5.**
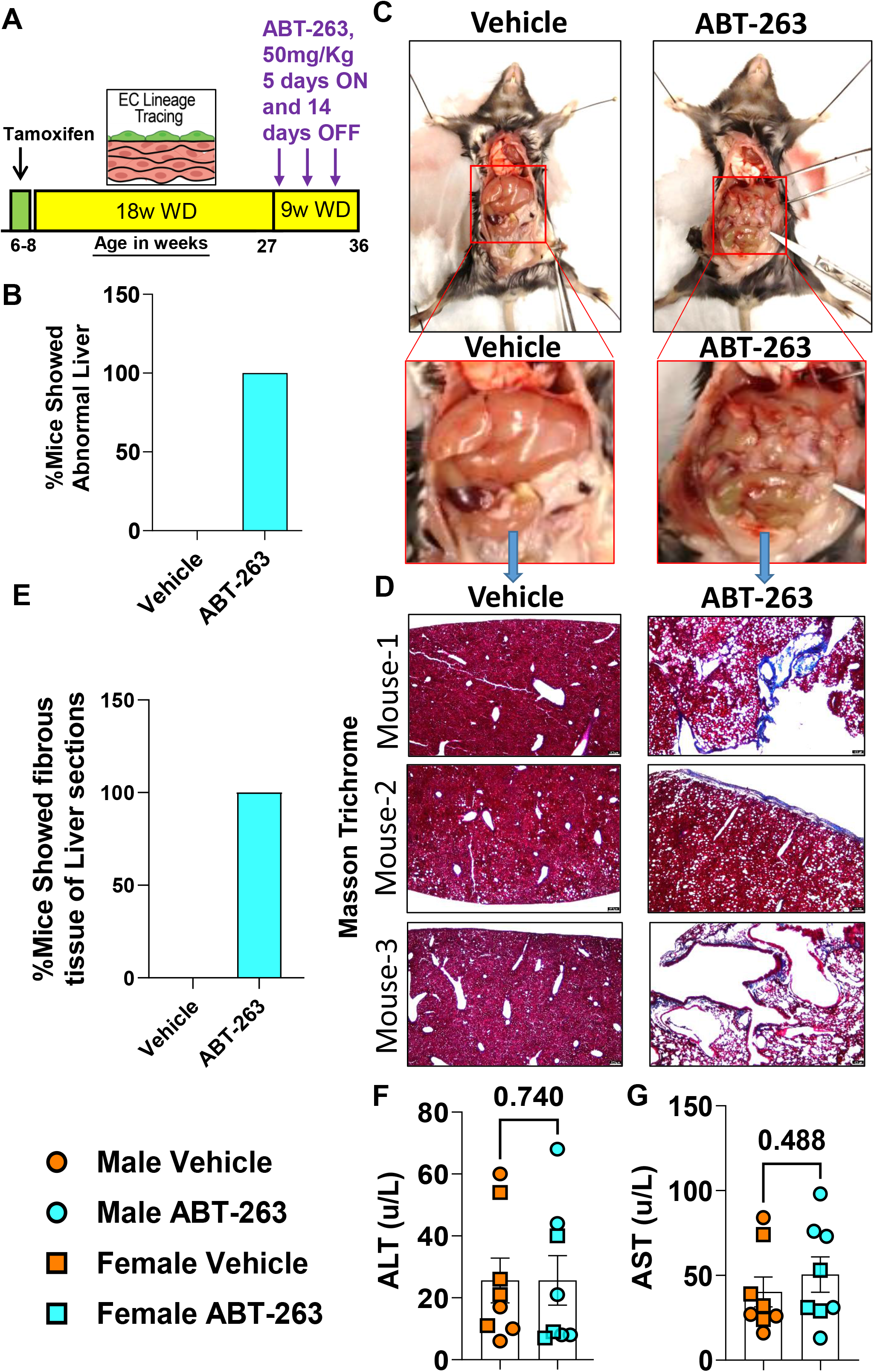
Low dose (50mg/kg/bw) ABT-263 treatment of *Apoe^-/-^* mice with advanced lesions was associated with increased hepatic fibrosis. (A) Experimental design, EC lineage tracing *Apoe*^-/-^ mice were fed a WD for 18 weeks followed by 50mg/kg/bw ABT 263 treatment on WD for 9 weeks. **(B)** Percentage of mice showing an abnormal liver phenotype. **(C)** Representative photographs of mice showing an abnormal live phenotype with ABT-263 treatment. **(D)** Masson trichrome stain to detect the fibrous tissue (blue) in the liver sections from the vehicle and ABT-263 treated mice. **(E)** Percentage mice showing fibrous tissue (blue) in the liver sections from the vehicle and ABT-263 treated mice. **(F)** ALT and **(G)** AST levels measured in plasma. Mann-Whitney U-tests were used for statistical analysis in F and G, and biologically independent animals are indicated as individual dots. Error bars show the SEM. The p-values are indicated on the respective graphs.

## Discussion

Senescent cells contribute to age-associated diseases^35^ and recent murine studies have tested the ability of senolytic agents to remove these senescent cells and to potentially treat a number of major diseases including atherosclerosis^25, 26, 28, 29^ neurodegeneration^36^, pulmonary fibrosis^37^, diabetic chronic kidney disease^38^, and cancer^39^. Indeed, multiple ongoing clinical trials with senolytic drugs are in progress targeting various diseases, including Alzheimer’s (NCT0463124), and diabetic chronic kidney disease (NCT02848131)^40^. These clinical trials are based on the belief that selective removal of these senescent cells will be beneficial. However, the results of the present study show that one of the most popular senolytic drugs, ABT-263, has multiple detrimental effects on *Apoe*^-/-^ mice with advanced atherosclerosis including a reduced probability of survival by 50-60%, that may be due to a reduced α-SMA^+^ fibrous cap thickness and a 90% reduction in SMC within the lesions. Indeed, our results suggest that the majority of SMC-derived cells within lesions, including those that are critical for the formation and maintenance of the protective fibrous cap, are particularly sensitive to ABT 263-induced clearance. Consistent with this possibility, Leeper and co-workers demonstrated that clonal expansion of lesion SMC is dependent on them escaping efferocytosis at least in part by activating the anti-phagocytic molecule CD47^41^. As such, results indicate that ABT-263 may be inducing apoptosis of non-senescent SMC and that with long-term use may increase the risk for the development of unstable advanced atherosclerotic lesions and the incidence of MI or stroke. This may be particularly severe since our data also showed that ABT-263 prevented adaptive increases in investment of EC-derived cells into the fibrous cap via beneficial EndoMT to myofibroblast transitions that we have shown normally occur when SMC investment into fibrous cap of lesions is impaired^9^.

At first glance, our results showing multiple detrimental changes in lesion pathology and a marked increase in mortality appear to be at odds with previous studies in the field reporting beneficial effects of ABT-263 treatment of atherosclerotic mice^25, 26^. The reasons for these differences are unclear, but likely include the following key variables between studies. **First,** to better match clinical paradigms of treating elderly patients with advanced disease, we did a late-stage intervention study^33^ involving initiation of ABT-263 treatment in *Apoe*^-/-^ mice after 18-weeks of WD (TD.88137; 42% calories from fat) feeding *so* that they already have highly advanced BCA lesions closely resembling advanced human lesions both morphologically^2^ and similar cellular composition of lesions as determined by scRNAseq analysis^15^. In contrast, previous studies showing beneficial effects of ABT-263 were either prevention studies where treatment was initiated in very young mice at the time of beginning WD feeding^25^, or an intervention study following just 12-14 weeks of WD feeding of *Ldlr*^-/-^ mice at which time lesions are still early stage^26^. **Second,** to model patients with undiagnosed advanced disease or with poor lipid management we elected to continue mice on a WD during the 9-week treatment period (i.e. 27 weeks of WD +/- ABT-263 treatment during the last 9 weeks). In contrast, to mimic the clinically relevant context of patients with highly effective medical management of pro-atherogenic lipids, Childs et al^26^ fed a WD to *Ldlr*^-/-^ mice for 12 weeks followed by 9 weeks of low fat diet (LFD) +/- ABT-263. They observed multiple beneficial effects of ABT-263 treatment including thickening of the fibrous cap of aortic lesions. Given that we modeled our intermittent ABT-263 dosing regimen to that of Childs et al., it is likely that the high mortality observed in our studies with ABT-263 is due, at least in part, to prolonged hyperlipidemia. However, it remains to be determined if our ABT-263-induced increase in mortality is due to thromboembolic events secondary to plaque de-stabilization, increased liver toxicity, and/or other mechanisms. Nevertheless, no matter what the cause of death with ABT-263 (***Navitoclax***), it is concerning given the extensive clinical^42, 43^ testing in progress. Indeed, results suggest that it may be prudent to exclude subjects with poorly controlled lipids, systemic inflammation, and/or advanced atherosclerosis from these studies.

Previous studies reported that EC-like^24^ and SMC-like^20^ cells undergo senescence that is associated with telomere shortening in human atherosclerotic lesions. The results of the present study showed that KO of *Klf4* in SMC was associated with reduced expression of pro-senescence markers, but preserved expression of the anti senescence marker, telomerase reverse transcriptase (*Tert*^+^). It is not clear how the loss of *Klf4* in SMC preserved *Tert*^+^ expression. However, it is probably the result of loss of *Klf4-*induced repression of *Tert* given previous studies showing that *Klf4* downregulates *hTERT* expression and telomerase activity to inhibit lung carcinoma growth^44^. Also the knockdown of *Klf4* in HUVECs in hyperglycemic conditions increased *hTERT* expression^45^. Telomerase maintains the telomere length and the telomerase deficiency (*Trf1*- and *Tert*-deficient mice) in mice lead to aplastic anemia and *Tert* gene therapy improved telomere length and blood count in these *Tert*-deficient mice^46^. Furthermore, recent *Tert* gene therapy in the mice, reduced vascular senescence and increased lifespan^47^, improved myocardial revascularization and tissue repair^48, 49^, and reduced neurodegeneration associated with short telomeres^50^. Given the multiple beneficial effects of *Tert* therapy in the pre-clinical mouse models, future studies need to evaluate if the *Tert* therapy would show beneficial effects on advanced atherosclerosis.

In conclusion, ABT-263 treatment of WD-fed *Apoe*^-/-^ mice with advanced atherosclerosis and persistent hyperlipidemia had multiple unexpected detrimental effects including it inducing reduced Myh11-eYFP^+^ SMC-derived cells within lesions and the fibrous cap and decreased α-SMA^+^ fibrous cap thickness. ABT-263 treatment also prevented adaptive increases in investment of EC-derived cells into the fibrous cap via beneficial EndoMT to myofibroblast transitions and increased mortality. Removing the senescent cells from the plaque is conceptually a good idea but given the multiple detrimental effects of ABT-263 in our WD fed *Apoe*^-/-^ mice, further pre-clinical studies with senolytic drugs are needed to identify factors and mechanisms that modulate their biological effects before planning a large clinical trial targeting atherosclerosis.

---

**Supplemental Figure I.**
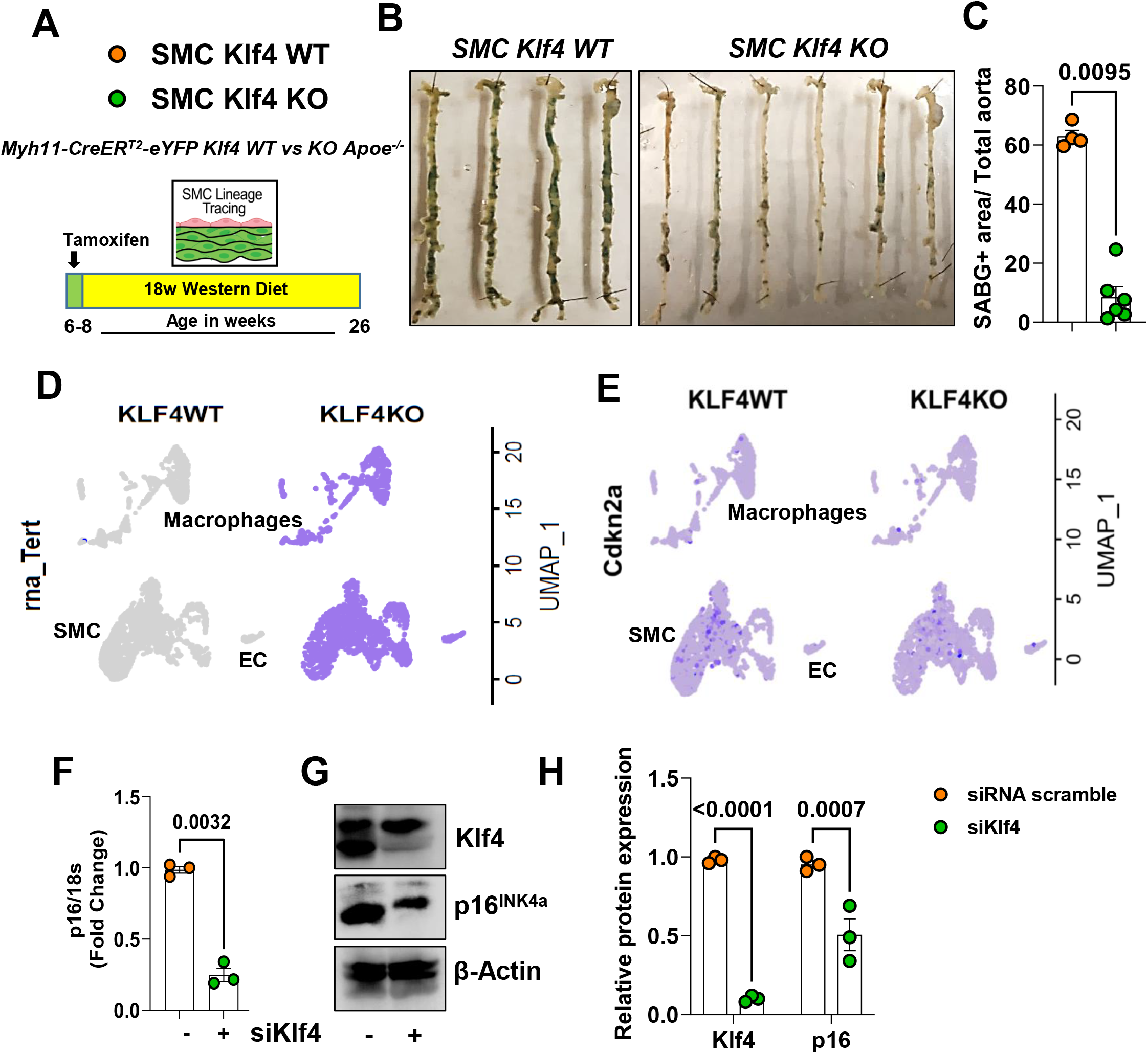
SMC Klf4 KO reduced overall lesion senescence. **(A)** Experimental design, SMC lineage tracing Klf4 WT vs KO *Apoe*^-/-^ mice were injected with tamoxifen at 6 to 8 weeks of age and subsequently placed on a western diet (WD) for 18 weeks to induce advanced atherosclerosis. Freshly isolated aortas were then **(B)** SAβG stained and **(C)** SAβG^+^ area normalized to total aorta shown in figure B quantified using Fiji (ImageJ 1.53c) software on digitized images. UMAPs of **(D)** *Tert* and **(E)** *p16* (*Cdkn2a*) from SMC Klf4 WT and KO scRNAseq data sets of BCA lesions (blue dots indicates the presence of cells). Cultured murine aortic SMC were transfected with siKlf4 for 24h and **(F)** *p16* mRNA levels were measured by qRT-PCR, and **(G)** p16 protein levels were measured by western blot, **(H)** Relative protein expressions of p16 and Klf4 were quantified with Fiji. Mann-Whitney U-tests were used for C, F and two-way ANOVA used for H. The error bars show the standard error of the mean (SEM). Independent animals are indicated as individual dots on the graphs. The p-values are indicated on the respective graphs.

**Supplemental Figure II.**
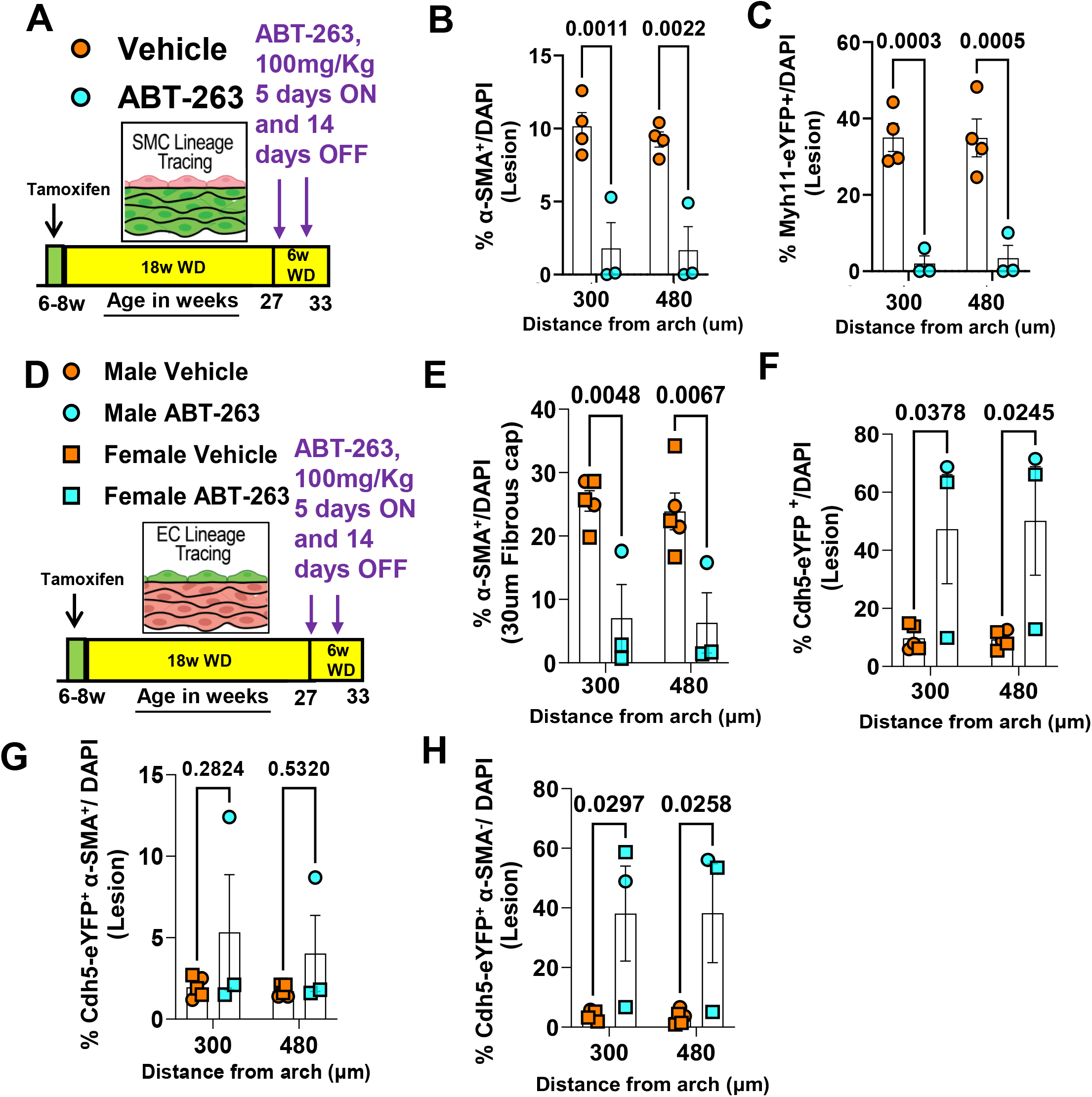
Treatment of SMC (Myh11-CreER^T2^-eYFP) and EC (Cdh5-CreER^T2^-eYFP)-lineage tracing *Apoe*^-/-^ mice with advanced atherosclerotic lesions with the senolytic drug ABT-263 (100mg/kg/bw) reduced the α-SMA^+^ cells but increased EC derived cells in BCA lesions. (A) Experimental design for B and C, SMC-lineage tracing *Apoe*^-/-^ mice were fed a WD for 18 weeks followed by ABT-263 treatment on a Western diet (WD) for 6 weeks. (B) α-SMA^+^ cells of all DAPI^+^ cells and (C) Myh11-eYFP^+^ (SMC) of all DAPI^+^ cells in the lesion. (D) Experimental design for figures E-H, EC-lineage tracing *Apoe*^-/-^ mice were fed a WD for 18 weeks followed by 100mg/kg/bw ABT-263 treatment on WD for 6 weeks. Note: Males and females data combined due to low n-number (Circles indicate males, and Squares indicate females). (E) Percentage of α-SMA^+^ cells in the fibrous cap of all (DAPI^+^) cells. (F) Percentage of endothelial cells (Cdh5-eYFP^+^) of all (DAPI^+^) cells in the lesions. (G) Percentage of EC-derived α-SMA^+^ (Cdh5-eYFP^+^ α-SMA^+^) cells of all (DAPI^+^) cells in the lesions. (H) Percentage of EC-derived but α- SMA negative (Cdh5-eYFP^+^ α-SMA^-^) cells of all (DAPI^+^) cells in the lesions. Two-way ANOVA used for statistics (B-C and E-I). The exact p-value is indicated on the respective graphs. The error bars show the standard error of the mean (SEM). Independent animals are indicated as individual dots on the graphs. The p-values are indicated on the respective graphs.

**Supplemental Figure III.**
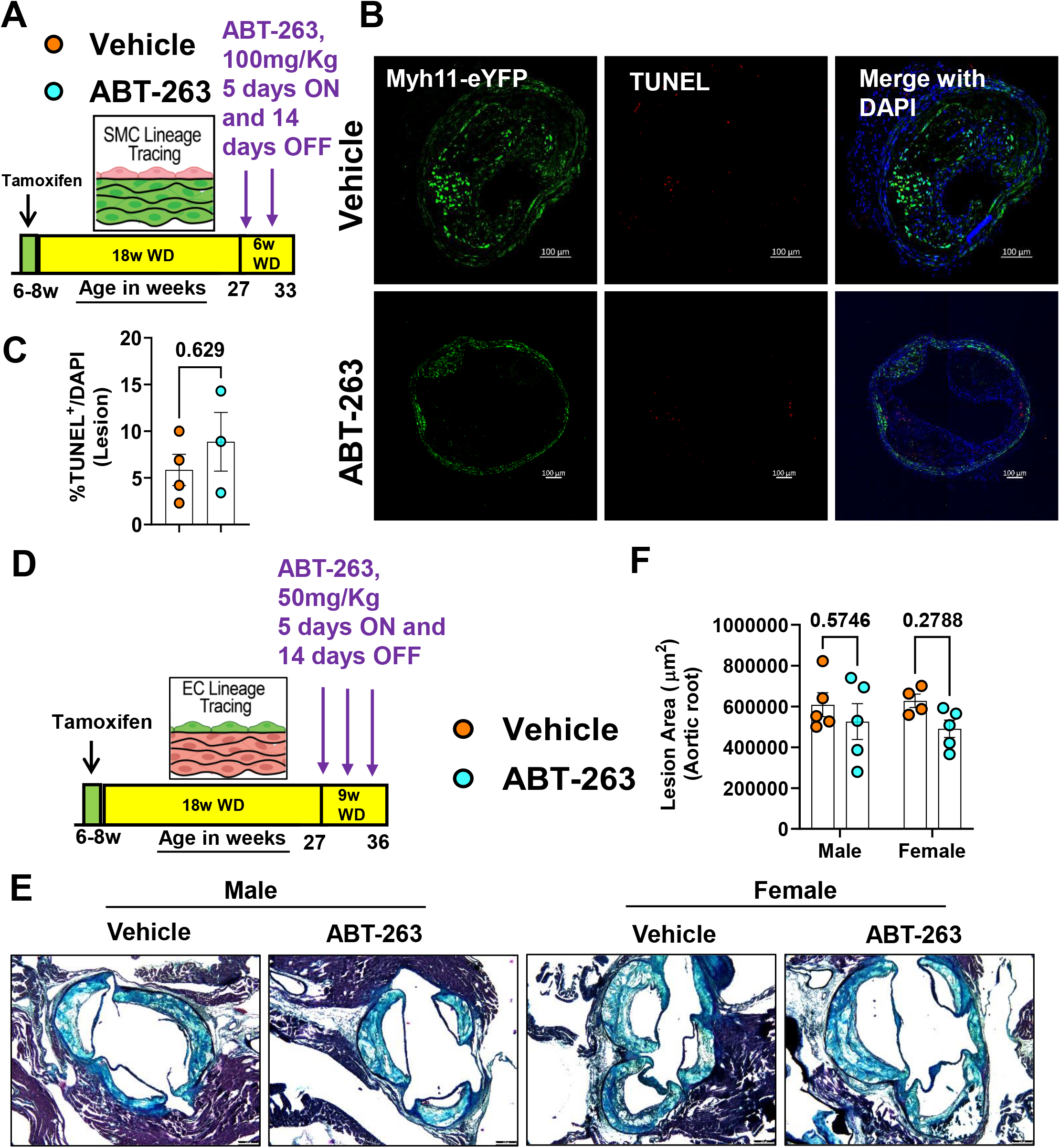
Treatment of *Apoe*^-/-^ mice with advanced atherosclerotic lesions with the senolytic drug ABT-263 (100mg/kg/bw) did not change the TUNEL^+^ apoptotic cells and a low dose (50mg/kg/bw) of ABT-263 treatment in *Apoe*^-/-^ mice with advanced lesions did not change the aortic root lesion size. (A) Experimental design for B and C, SMC-lineage tracing *Apoe*^-/-^ mice were fed a WD for 18 weeks followed by ABT-263 treatment on a Western diet (WD) for 6 weeks. (B) Myh11-eYFP (Green), TUNEL (Red), and Merged with DAPI on BCA lesions (C) TUNEL+ of all DAPI+ cells in the lesion. (D) Experimental design, EC-lineage tracing *Apoe*^-/-^ mice were fed a WD for 18 weeks followed by 50mg/kg/bw ABT-263 treatment on WD for 9 weeks. (E) Representative 10x images with 100μm scale bar of MOVAT staining on Aortic root lesions. (F) Quantification of lesion area from E. Mann-Whitney U-tests used for C and Two-way ANOVA test used for F. The error bars show the standard error of the mean (SEM). Independent animals are indicated as individual dots on the graphs. The p-values are indicated on the respective graphs.

**Supplemental Figure IV.**
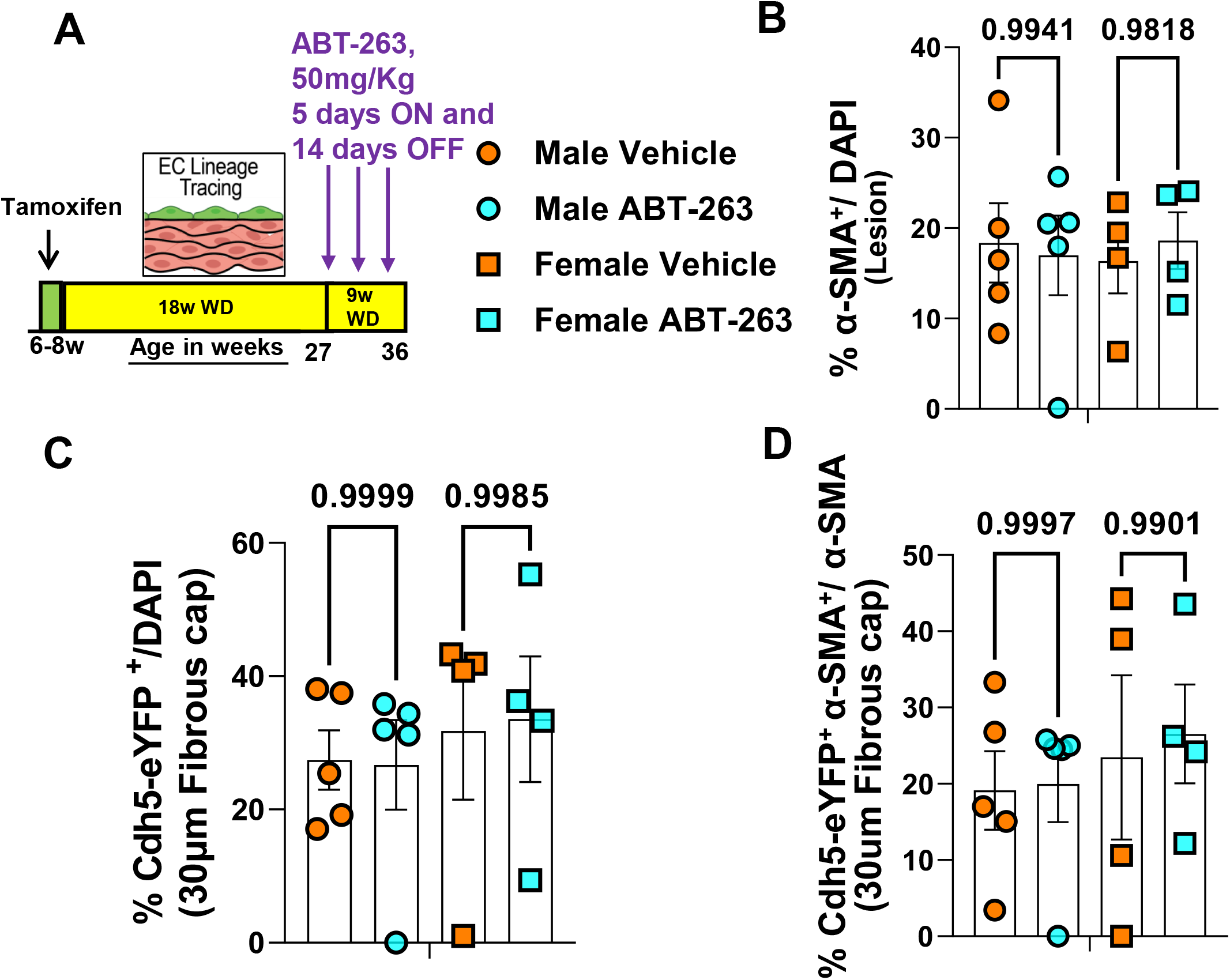
A reduced dose (50mg/kg/bw) of ABT-263 treatment on *Apoe*^-/-^ mice with advanced lesions inhibited investment of EC-derived cells into the fibrous cap via beneficial EndoMT to myofibroblast transitions. (A) Experimental design, EC-lineage tracing *Apoe*^-/-^ mice were fed a WD for 18 weeks followed by 50mg/kg/bw ABT-263 treatment on WD for 9 weeks. (B) α-SMA^+^ cells of all (DAPI^+^) cells in the lesion. (C) Endothelial cells of all (DAPI^+^) cells in the fibrous cap. (D) EC-derived α-SMA^+^ cells of all α-SMA^+^ cells in the fibrous cap. Two-way ANOVA used for statistics. The error bars show the standard error of the mean (SEM). Independent animals are indicated as individual dots on the graphs. The p-values are indicated on the respective graphs.

**Supplemental Figure V.**
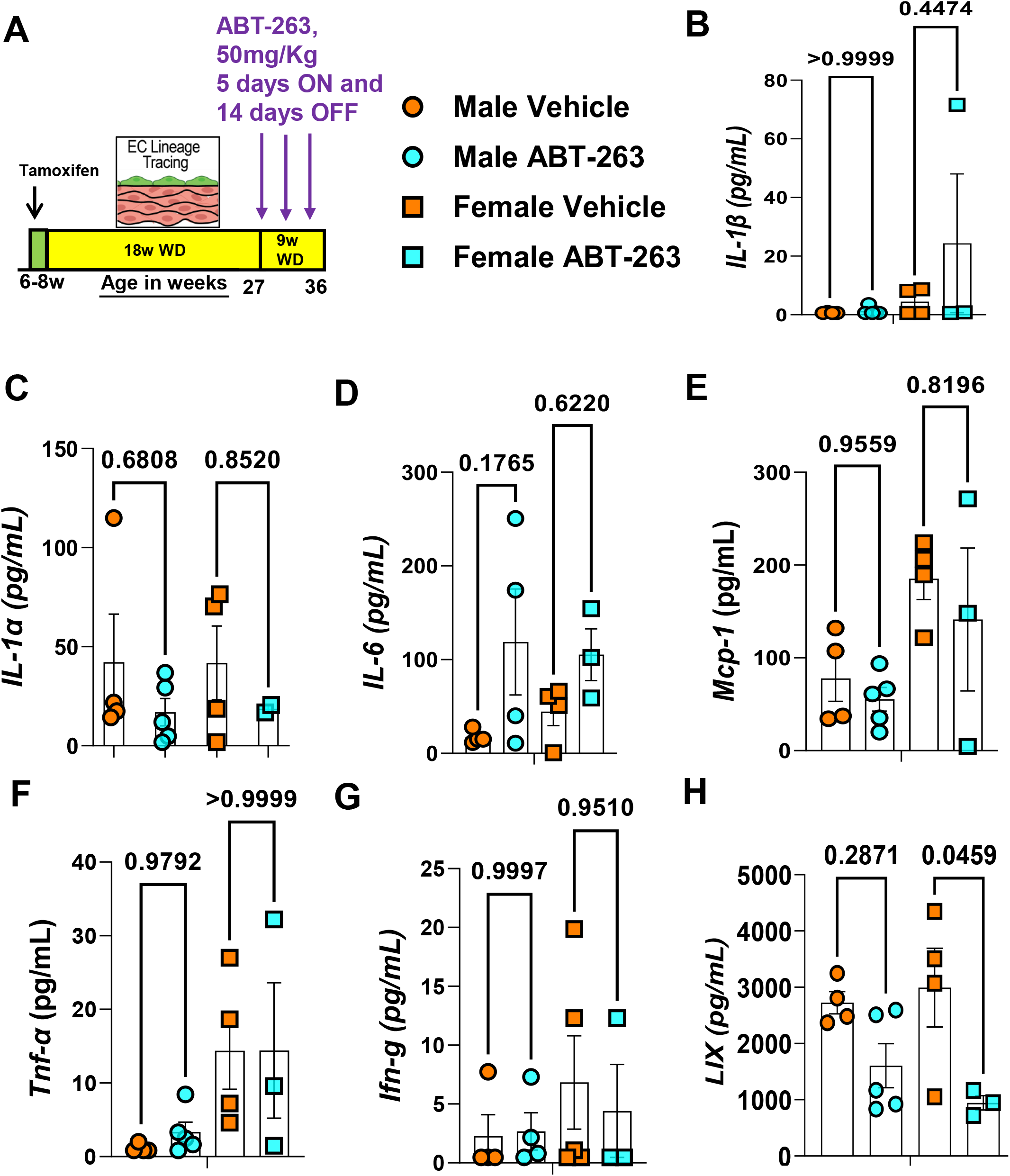
Low dose ABT-263 treatment of WD-fed *Apoe*^-/-^ mice reduced plasma Cxcl5, but did not reduce SASPs and cytokines including, IL-1β, IL-1α, IL-6, Mcp-1, Tnf-α, and Ifn-g. (A) Experimental design, EC-lineage tracing *Apoe*^-/-^ mice were fed a WD for 18 weeks followed by 50mg/kg/bw ABT-263 treatment on WD for 9 (3 cycles 5 days ON and 14 days OFF) weeks. Plasma (B) IL-1β, (C) IL-1α, (D) IL-6, (E) Mcp-1, (F) Tnf-α, (G) Ifn-g, and LIX (Cxcl5) (H). Two-way ANOVA used for statistics. The error bars show the standard error of the mean (SEM). Independent animals are indicated as individual dots on the graphs. The p-values are indicated on the respective graphs.

**Supplemental Figure VI.**
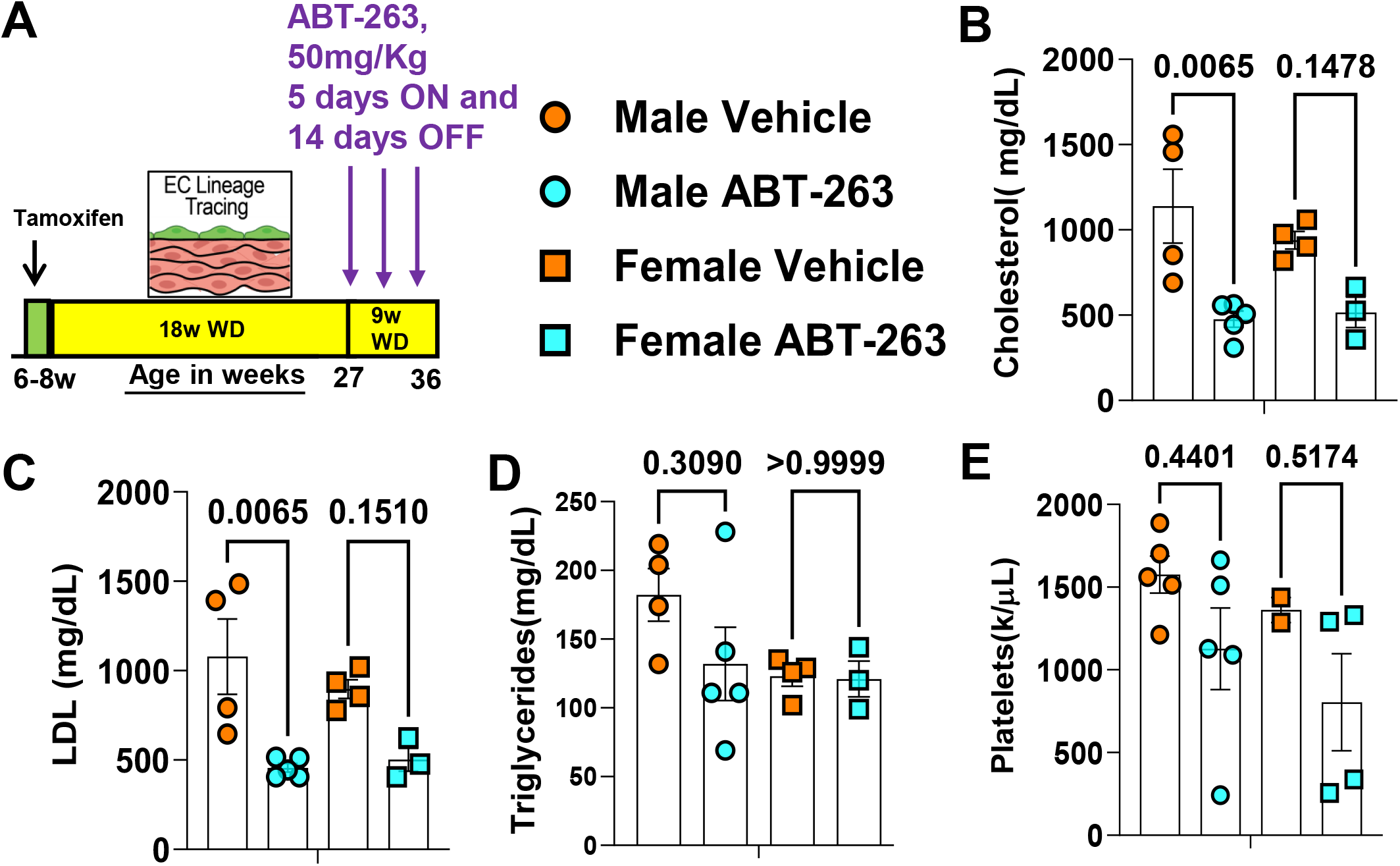
Low dose ABT-263 treatment of WD-fed *Apoe*^-/-^ mice reduced plasma lipids. **(A)** Experimental design, EC-lineage tracing *Apoe*^-/-^ mice were fed a WD for 18 weeks followed by 50mg/kg/bw ABT-263 treatment on WD for 9 weeks. **(B)** Plasma cholesterol, LDL **(C)** LDL, **(D)** Triglycerides, and **(E)** Platelets in the blood. Two-way ANOVA used for statistics (B-E), biologically independent animals are indicated as individual dots, and error bars show SEM. The exact p-value is indicated on the respective graph. The exact p-value is indicated on the respective graph.

**Supplemental Figure VII.**
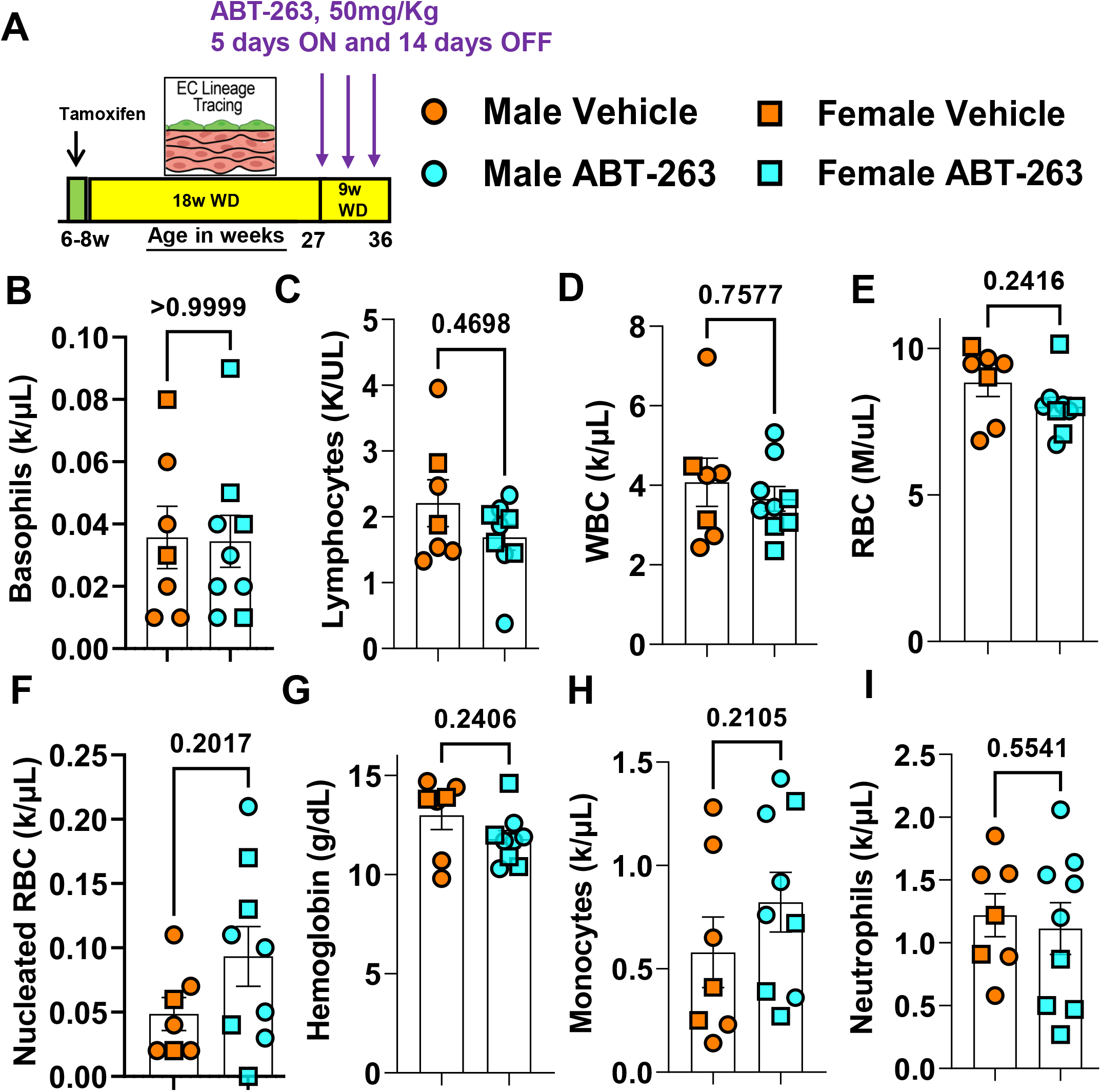
Low dose ABT-263 treatment of WD-fed *Apoe*^-/-^ mice reduced plasma lipids. **(A)** Experimental design, EC-lineage tracing *Apoe*^-/-^ mice were fed a WD for 18 weeks followed by 50mg/kg/bw ABT-263 treatment on WD for 9 weeks. **(B)** Basophils, **(C)** Lymphocytes, **(D)** WBC, **(E)** RBC, **(F)** Nucleated RBC, **(G)** Hemoglobin, **(H)** Monocytes, and **(I)** Neutrophils in the blood. Mann-Whitney U-tests used for statistics. The error bars show the standard error of the mean (SEM). Independent animals are indicated as individual dots on the graphs. The p-values are indicated on the respective graphs.

**Supplemental Figure VIII.**
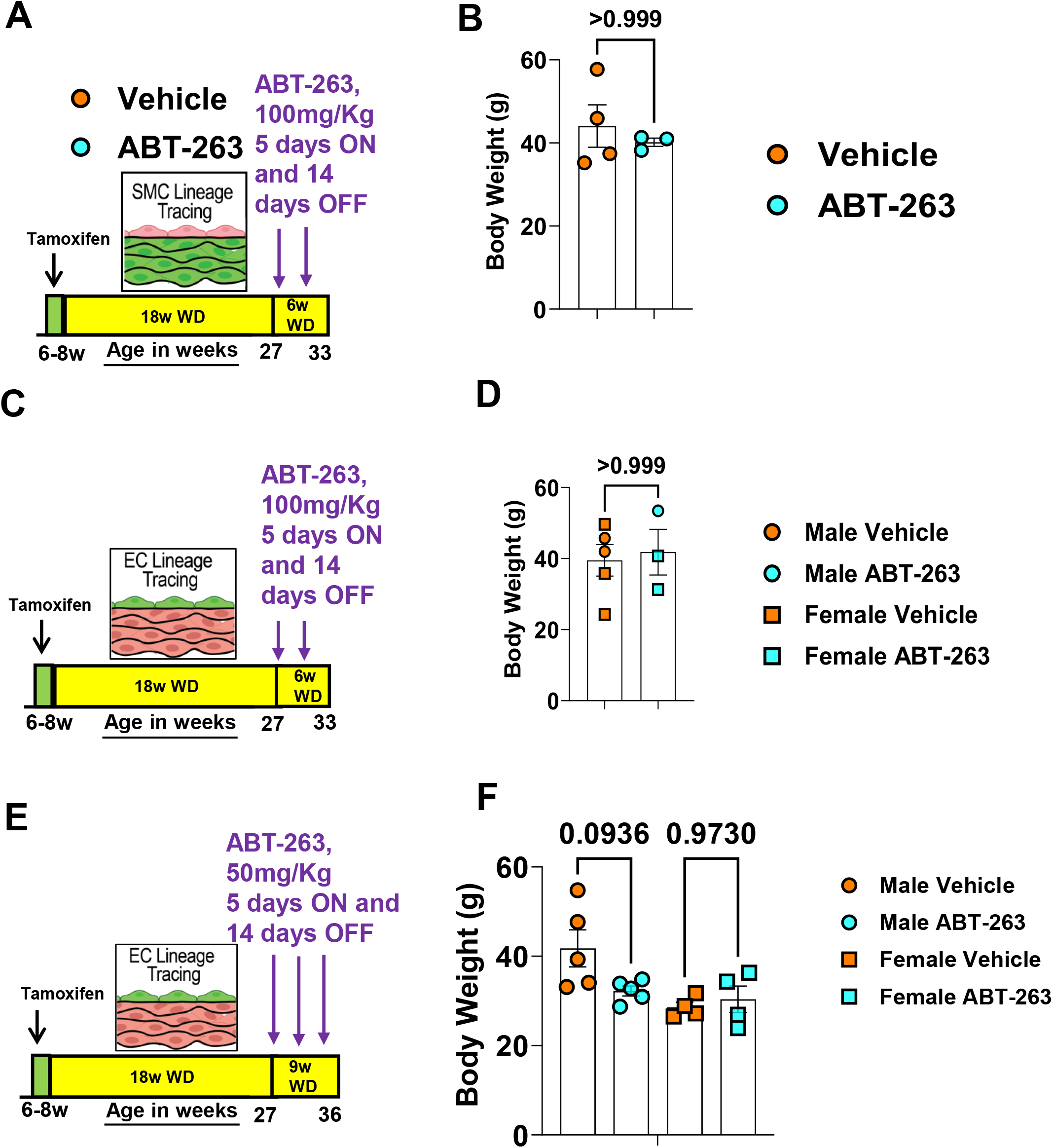
ABT-263 treatment of WD-fed *Apoe*^-/-^ mice did not affect the body weight. **(A)** Experimental design, SMC-lineage tracing *Apoe*^-/-^ mice were fed a WD for 18 weeks followed by 100mg/kg/bw ABT-263 treatment on WD for 6 weeks. **(B)** Body weight (g), **(C)** Experimental design, EC-lineage tracing *Apoe*^-/-^ mice were fed a WD for 18 weeks followed by 100mg/kg/bw ABT-263 treatment on WD for 6 weeks. **(D)** Body weight (g), **(E)** Experimental design, EC-lineage tracing *Apoe*^-/-^ mice were fed a WD for 18 weeks followed by 50mg/kg/bw ABT-263 treatment on WD for 9 weeks. **(F)** Body weight (g). Mann-Whitney U-tests used for B and D, Two-way ANOVA used for F. The error bars show the standard error of the mean (SEM). Independent animals are indicated as individual dots on the graphs. The p-values are indicated on the respective graphs.

